# Genetic rescue of pathogenic O-GlcNAc dyshomeostasis associated with microcephaly and motor deficits

**DOI:** 10.1101/2025.11.12.687959

**Authors:** Florence Authier, Iria Esperon-Abril, Kévin-Sébastien Coquelin, Christian Stald Skoven, Simon Fristed Eskildsen, Nina Ondruskova, Andrew T. Ferenbach, Jesper Skovhus Thomsen, Brian Hansen, Daan M. F. van Aalten

## Abstract

Missense variants in O-GlcNAc transferase (OGT) result in OGT congenital disorder of glycosylation (OGT-CDG), an intellectual disability syndrome associated with O-GlcNAc dyshomeostasis and a range of neurodevelopmental defects. Inhibition of O-GlcNAcase (OGA), the enzyme responsible for removing protein O-GlcNAcylation, has been explored as a target for modulating brain O-GlcNAc homeostasis in neurodegenerative diseases and may also be a target for OGT-CDG. Here, we describe an OGT-CDG mouse line that exhibits microcephaly, motor deficits, and brain O-GlcNAc dyshomeostasis, closely mirroring patient symptoms. We genetically explored OGA as a target for OGT-CDG by crossing these mice with a line carrying catalytically inactive OGA. Encouragingly, this partially restored O-GlcNAc homeostasis in brain and blood, although it did not result in significant phenotypic rescue. These findings suggest that OGA inhibition can modulate enzymatic imbalance in OGT-CDG mice, and that blood can be used to monitor the effects of interventions targeting O-GlcNAc dyshomeostasis.

## Introduction

The maintenance of cellular homeostasis is crucial for proper cell function and requires mechanisms that can respond rapidly to metabolic and environmental changes. This regulation is primarily controlled by post-translational modifications (PTMs), which lead to rapid, reversible changes in protein dynamics. One such metabolism-responsive PTM is O-linked β-N-acetylglucosamine (O-GlcNAc) (1,2). O-GlcNAc moieties are covalently attached to serine and threonine residues on thousands of proteins localized in the nucleus, cytoplasm, and mitochondria (3,4). Unlike many other PTMs, O-GlcNAc cycling is controlled by only two enzymes: O-GlcNAc transferase (OGT), which adds the modification, and O-GlcNAcase (OGA), which removes it (5,6). O-GlcNAc integrates cellular nutrient availability and metabolic state, and its levels depend on the tight regulation of both OGT and OGA (7–10). Accordingly, protein O-GlcNAcylation is implicated in the regulation of numerous crucial cellular processes, including gene expression, stress responses, and metabolic pathways (10–12) and is linked to a range of diseases (13,14).

O-GlcNAcylation and its associated enzymes play a crucial role in normal development, as shown by the lethality of knockout animal models (15–18). The enzymes and the modification are particularly abundant in the brain (19–21), with several studies indicating that O-GlcNAc plays a role in neurodevelopment and synaptic function (22–26). Accordingly, the levels of O-GlcNAc, OGT, and OGA are tightly regulated during mammalian brain development, showing a gradual decline of global O-GlcNAcylation with age (22). Disrupted O-GlcNAc levels have been associated with several neurodegenerative diseases, including Parkinson’s and Alzheimer’s disease (AD) (27,28). AD is characterized, among other features, by reduced O-GlcNAc levels in the brain, which have been linked to increased necroptosis, neuroinflammation, and tau pathology (29). O-GlcNAcylation of tau anti-correlates with pathogenic hyperphosphorylation, suggesting that the reduction in O-GlcNAc levels in AD could contribute to tauopathies (29–31). In mouse models of AD, increasing O-GlcNAc by chronic treatment with OGA inhibitors had a neuroprotective effect against both tau and amyloid-β peptide pathology (29,32–34), although most of these inhibitors have now failed in the clinic due to neurotoxicity (35). Similar conclusions were drawn when examining the impact of increasing O-GlcNAc on Parkinson’s disease pathology (36–39), epilepsy (40,41), ALS (42–44), and Down syndrome (45,46). In contrast, other studies indicate that OGA inhibition is detrimental to neurons via α-synuclein accumulation (47) and that decreased O-GlcNAcylation has a protective effect in animal models of other neurodegenerative disorders (48,49). Overall, the mechanisms mediating the impact of O-GlcNAc in neurodegenerative diseases are not well understood, as changes in O-GlcNAc homeostasis may have context-and disease-dependent impacts.

Recently, pathogenic missense variants in the *OGT* gene have been linked to a syndromic form of X-linked intellectual disability (ID), known as O-GlcNAc Transferase Congenital Disorder of Glycosylation (OGT-CDG) (50). The OGT-CDG syndrome is characterised by several neurological and psychiatric manifestations, such as developmental delay, autism spectrum disorder (ASD), and epilepsy, as well as locomotion defects and behavioural deficits. Several OGT-CDG variants have been clinically described (51–56), revealing that patients typically exhibit a range of additional facial and skeletal abnormalities, including microcephaly, craniofacial dysmorphia, clinodactyly, and growth impairment (50,57). There are no treatments available for this group of patients, and it is not yet known whether the neurological symptoms are primarily of neurodevelopmental or neurophysiological origin. When modelled in stem cells, *Drosophila* and mice, these missense variants cause O-GlcNAc dyshomeostasis, presumably as a result of reduced OGT activity, and it is assumed that loss of O-GlcNAc on key proteins contributes to the neurological phenotypes observed, through mechanisms that remain to be dissected (51,56–60).

In OGT-CDG, several characterised variants result in a decrease in OGA protein levels (51,52,56), likely exploited as a compensatory mechanism by cells to attempt to restore O-GlcNAc homeostasis. This would suggest that inhibiting OGA in the context of OGT-CDG may be a potential method for (further) correcting O-GlcNAc imbalance. However, it is not clear whether such OGA inhibition would be required during early development to compensate for neurodevelopmental defects during embryogenesis and/or in adulthood to compensate for neurophysiological defects. We sought a genetic strategy to dissect these possible mechanisms and explore OGA as a target for OGT-CDG. We have previously reported an OGA-catalytically deficient mouse (*Oga*^D285A^) generated through a constitutive knock-in of an Asp285Ala mutation at an active site residue that is key for both catalytic activity and O-GlcNAc binding (15). Homozygous *Oga*^D285A/D285A^ mice, like the previously reported *Oga* knockout mice (17), die perinatally, emphasizing the crucial role of OGA activity during development. However, heterozygous *Oga*^D285A/+^ mice develop normally and exhibit tissue-specific changes in O-GlcNAc homeostasis regulation to compensate for OGA catalytic deficiency (15). Here, we used these mice to restore O-GlcNAc homeostasis in a mouse model of OGT-CDG. To achieve this, we first generated a new OGT-CDG mouse line carrying the N648Y patient variant (*Ogt*^N648Y/y^). This *de novo* variant leads to the non-conservative substitution of Asn648 in the OGT active site with Tyr, resulting in a significant loss of activity of the enzyme, hypo-O-GlcNAcylation in a stem cell model, and significant neurodevelopmental deficits in the patient (51). *Ogt*^N648Y/y^ mice exhibit a reduction in brain, skull, and body size, brain and blood O-GlcNAc levels, and behavioural defects that recapitulate symptoms seen in the patient, such as hyperactivity and motor deficits. We generated *Ogt*^N648Y/y^*Oga*^D285A/+^ mice, lacking 50% of OGA activity from embryogenesis to adulthood, as a genetic approach to compensate for the hypo-O-GlcNAcylation seen in the *Ogt*^N648Y/y^ mice. Although this did not lead to improvements in OGT-CDG-related behavioural deficits or body weight, we found that this approach did partially restore O-GlcNAc homeostasis in the brain and blood, emphasizing the potential of blood samples as a disease biomarker and evaluating the efficiency of future treatments for OGT-CDG.

## Results

### Ogt^N648Y/y^ mice are viable and exhibit brain O-GlcNAc dyshomeostasis

To investigate the effects of OGT-CDG variants *in vivo*, we focused on a recently described missense variant, N648Y, the first OGT-CDG variant reported within the catalytic domain of OGT (51). This variant was identified *de novo* in the second child of a non-consanguineous family, where neither parent nor the older brother carried the variant and were all healthy. Facial asymmetry was evident shortly after birth, while other symptoms of the disease were noticeable only a few months later, as he showed a delay in achieving critical developmental milestones. In addition to physical features such as syndactyly and persistent drooling, the patient had no speech and presented moderate hyperactivity and increased sensitivity to external stimuli during early childhood. Given the severity of the clinical manifestations, the N648Y variant became a strong candidate for *in vivo* modelling. We used a CRISPR/Cas9 genome editing approach in mice to introduce this variant, generating cohorts of *Ogt*^N648Y/y^ males for phenotypic analyses (**Fig. 1A**). Briefly, pseudo-pregnant female mice were implanted with 0.5 days post-coitum zygotes injected with editing reagents. DNA from offspring was genotyped and sequenced to confirm the presence of the OGT^N648Y^ variant (**Fig. 1B**). As the gene encoding *OGT* is located on the X chromosome in both mouse and human, we evaluated the Mendelian inheritance distribution of male offspring generated from heterozygous *Ogt*^N648Y/+^ females crossed to wild-type (WT) *Ogt*^+/y^ males. We observed that the crosses generated 58 % of male *Ogt*^+/y^ and 42 % of male *Ogt*^N648Y/y^ animals, indicating a slightly reduced percentage of *Ogt*^N648Y/y^ animals born (**Fig. 1C**). We observed that *Ogt*^N648Y/y^ mice exhibit a decrease in body (**Fig. 1D**) and brain (**Fig. 1E**) weight compared to their WT littermates similar to that described in the recent first report of an OGT-CDG mouse model (56,61). Taken together, these results indicate that mice carrying the OGT^N648Y^ variant are viable but show reduced growth. Given the X-linked inheritance of *Ogt*, only WT and hemizygous *Ogt*^N648Y/y^ male animals were used for phenotypic analyses.

**Figure 1:**
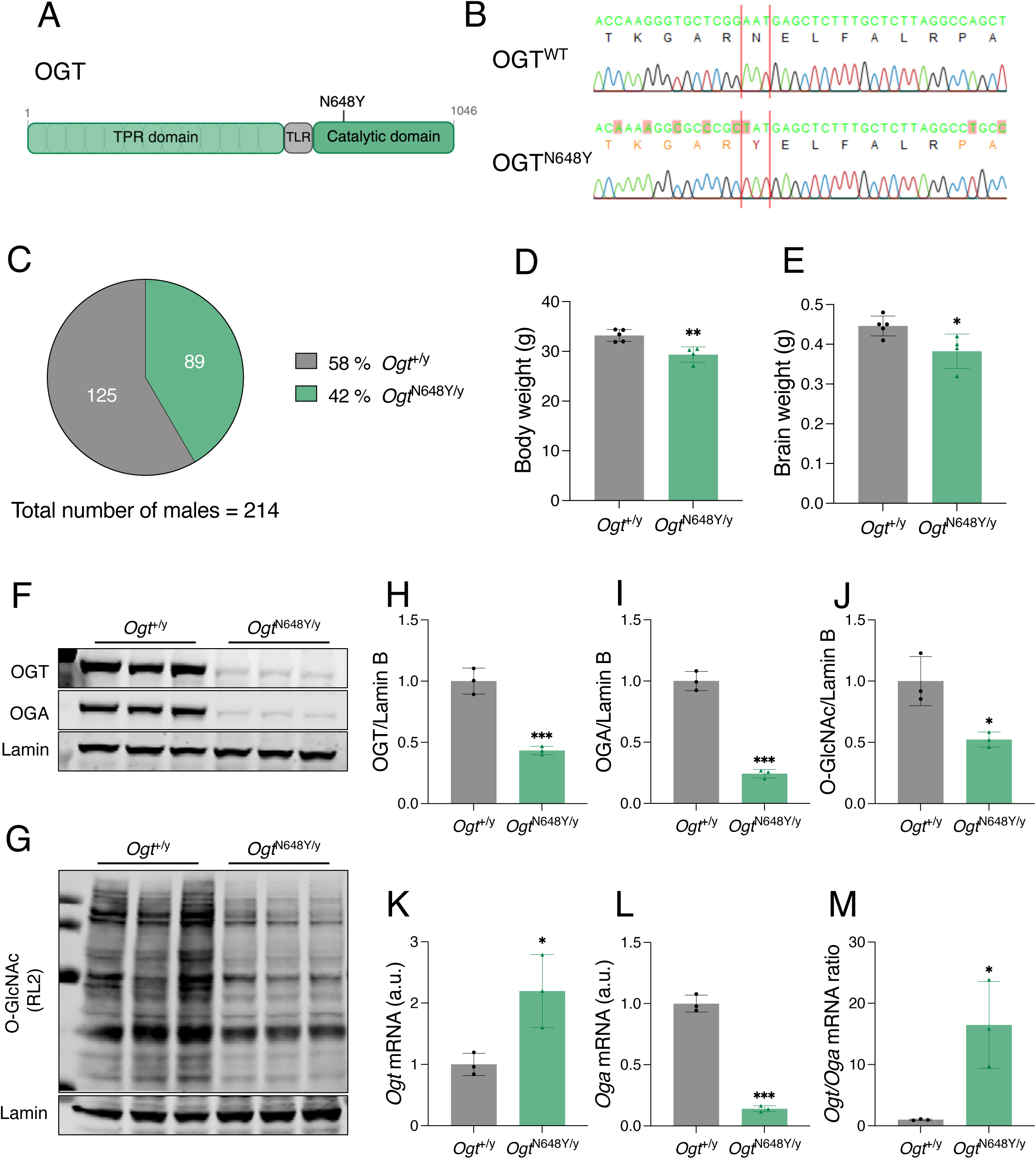
OGT^N648Y^ variant causes O-GlcNAc dyshomeostasis in the mouse brain. Data are represented as mean ± SD. Student *t*-test is used for statistics. Significance is shown as: \**p* < 0.05, \*\**p* < 0.01 and \*\*\**p* < 0.001 **(a)** Schematic of the OGT protein with the OGT^N648Y^ variant. The TPR (light green), TLR (grey), and catalytic domain (dark green) of OGT are represented. **(b)** Sequencing of genomic DNA of OGT^WT^ (top) and OGT^N648Y^ (bottom) confirms the presence of the N648Y point mutation (highlighted in red) in the transgenic animals **(c)** Diagram showing numbers and percentages of *Ogt*^+/y^ and *Ogt*^N648Y/y^ animals generated from female *Ogt*^N648Y/+^ x male *Ogt*^+/y^ breeding pairs. **(d)** Measurement of body weight of *Ogt*^+/y^ (n = 5) and *Ogt*^N648Y/y^ (n = 4) mice. **(e)** Measurement of brain weight of *Ogt*^+/y^ (n = 5) and *Ogt*^N648Y/y^ (n = 4) mice. **(f)** Western blot of OGT and OGA levels in adult brain of *Ogt*^+/y^ (n = 3) and *Ogt*^N648Y/y^ (n = 3) mice. Lamin antibody was used as a loading control. **(g)** Western blot of O-GlcNAc levels in adult brain of *Ogt*^+/y^ (n = 3) and *Ogt*^N648Y/y^ (n = 3) mice. Lamin antibody was used as a loading control. **(h)** Quantification of OGT protein levels from the Western blot in panel (f). **(i)** Quantification of OGA protein levels from the Western blot in panel (f). **(j)** Quantification of O-GlcNAcylated proteins levels from the Western blot in panel (g). **(k)** Quantification of *Ogt* mRNA levels in whole mouse brain of *Ogt*^+/y^ (n = 3) and *Ogt*^N648Y/y^ (n = 3) by RT-PCR. **(l)** Quantification of *Oga* mRNA levels in whole mouse brain of *Ogt*^+/y^ (n = 3) and *Ogt*^N648Y/y^ (n = 3) by RT-PCR. **(m)** Quantification of *Ogt/Oga* mRNA ratio in whole mouse brain of *Ogt*^+/y^ (n = 3) and *Ogt*^N648Y/y^ (n = 3) by RT-PCR.

The OGT^N648Y^ variant was shown to affect OGT catalytic activity, leading to a reduction in OGA levels and hypoglycosylation in stem cells (51). Therefore, we assessed the levels of OGT, OGA, and O-GlcNAcylation in whole brains from adult WT and *Ogt*^N648Y/y^ animals. We observed that both OGT (**Fig. 1F, H**) and OGA (**Fig. 1F, I**) protein levels were significantly reduced in *Ogt*^N648Y/y^ mice compared to WT littermates. Likewise, global O-GlcNAcylated proteins were also significantly decreased in the *Ogt*^N648Y/y^ mice compared to WT (**Fig. 1G, J**). To determine whether the differences in OGT and OGA protein levels were due to impaired transcription, we performed qPCR analysis on whole-brain extracts. We found a significant increase in *Ogt* mRNA expression in *Ogt*^N648Y/y^ mouse brains (**Fig. 1K**), while *Oga* mRNA expression was decreased (**Fig. 1L**) compared to WT animals, leading to an increase in *Ogt/Oga* mRNA in *Ogt*^N648Y/y^ mice compared to WT – a sensitive measurement of O-GlcNAc dyshomeostasis (**Fig. 1M**). This indicates that the lower OGT protein levels seen in the *Ogt*^N648Y/y^ mutant are not caused by decreased *Ogt* mRNA expression but instead result from a post-translational process. Taken together, these data suggest that while *Ogt*^N648Y/y^ mice are viable, they exhibit brain O-GlcNAc dyshomeostasis.

### Ogt^N648Y/y^ mice exhibit microcephaly associated with cortical hypoplasia

To assess signs of global developmental delay, body weight was monitored weekly (**Fig. 2A**). In the first two weeks after weaning (4 to 5 weeks old), *Ogt*^N648Y/y^ and WT mice showed comparable body weights, whereas after 5 weeks old, lower gains in body weights in *Ogt*^N648Y/y^ mice were observed compared to their WT littermates (**Fig. 2A**), suggesting that *Ogt*^N648Y/y^ mice show general delayed postnatal development. OGT-CDG patients exhibit craniofacial abnormalities, including microcephaly, also seen in the patient carrying the OGT^N648Y^ variant (51). Therefore, we performed a comprehensive analysis of skull and brain morphometrics. We assessed skull shape and size using micro-computed tomography (µCT) imaging in *Ogt*^N648Y/y^ and WT mice at 8 weeks (WT, n = 5; *Ogt*^N648Y/y^, n = 8) and 20 weeks of age (WT, n = 11; *Ogt*^N648Y/y^, n = 12). For each animal, models were generated of the skull and the extracted endocranial cavity (endocast) as a proxy for brain volume (**Fig. 2B**). Measurement of the skull volume indicated that 20-week-old *Ogt*^N648Y/y^ mice show a significantly lower bone volume compared to their WT littermates. Similarly, 8-week-old *Ogt*^N648Y/y^ mice showed lower skull volume compared to WT animals, albeit not significant (**Fig. 2C**). Next, we measured the volume of the endocast as a proxy for brain volume. *Ogt*^N648Y/y^ mice exhibited 15 % smaller endocasts than their WT littermates at both ages (**Fig. 2D**). Interestingly, the skull (bone) volume of mice from each genotype increased between 8 and 20 weeks of age, while the endocast volume remained unchanged (**Fig. 2C, D**), indicating that the bone continued to grow independently of the brain. To investigate whether the lower skull volume in the *Ogt*^N648Y/y^ mice was due to global or regional differences in size, we performed Euclidean Distance Matrix Analysis (EDMA) between 45 skull surface markers (61). At 20 weeks of age, 56 % of the surface distances in *Ogt*^N648Y/y^ mouse skulls were at least 2 % shorter compared to WT, with reductions of at least 5 % observed in both the anteroposterior and vertical (cranial height) dimensions (**Fig. 2E**). To identify differences in skull shape independently of size differences, we used surface landmarks on the skulls to perform Principal Components Analysis (PCA) in the 20-week-old mouse skulls (**Fig. 2F**). We observed that, while most of the WT mouse skulls separated along the first and second principal components (PC1 and PC2, respectively), *Ogt*^N648Y/y^ mouse skulls appeared more dispersed, indicating a higher morphological heterogeneity. Next, we turned to magnetic resonance imaging (MRI) to determine whether the reduction in brain size in the 20-week-old *Ogt*^N648Y/y^ mice is global or localized to specific brain regions. As expected, the *Ogt*^N648Y/y^ mice were found to have significantly smaller total brain volumes compared to WT littermates (**Fig. 2G**). Regional volumetric analysis showed a reduction in absolute volumes in most brain regions (**Fig. S1**). Regional relative volumes (normalized to total brain volume) in 39 bilaterally pooled brain regions were also determined (**Fig. S2**). The mean and median relative volumes were significantly different between the groups in the following brain regions: anterior commissure, cerebellar peduncle, inferior colliculus, fimbria, lateral septum, and olfactory bulbs. Considering only the means, differences were found in the corpus callosum and olfactory tubercle; while when only considering the medians, the areas that were significantly different were the midbrain and the periaqueductal grey. However, none of the differences in the regional relative volumes was significant when correcting for multiple comparisons (penalizing for the number of areas and Benjamini-Hochberg). Cortical thickness analysis revealed significantly thinner cortex over both hemispheres in the *Ogt*^N648Y/y^ mice compared to WT, with this difference remaining significant even after correcting for family-wise error (FWE) (**Fig. 2H**). Taken together, these results suggest that the *Ogt*^N648Y/y^ mice exhibit a reduced skull size and shape deformations compared to their WT littermates, suggesting microcephaly and changes in skull shape, associated with cortical hypoplasia.

**Figure 2:**
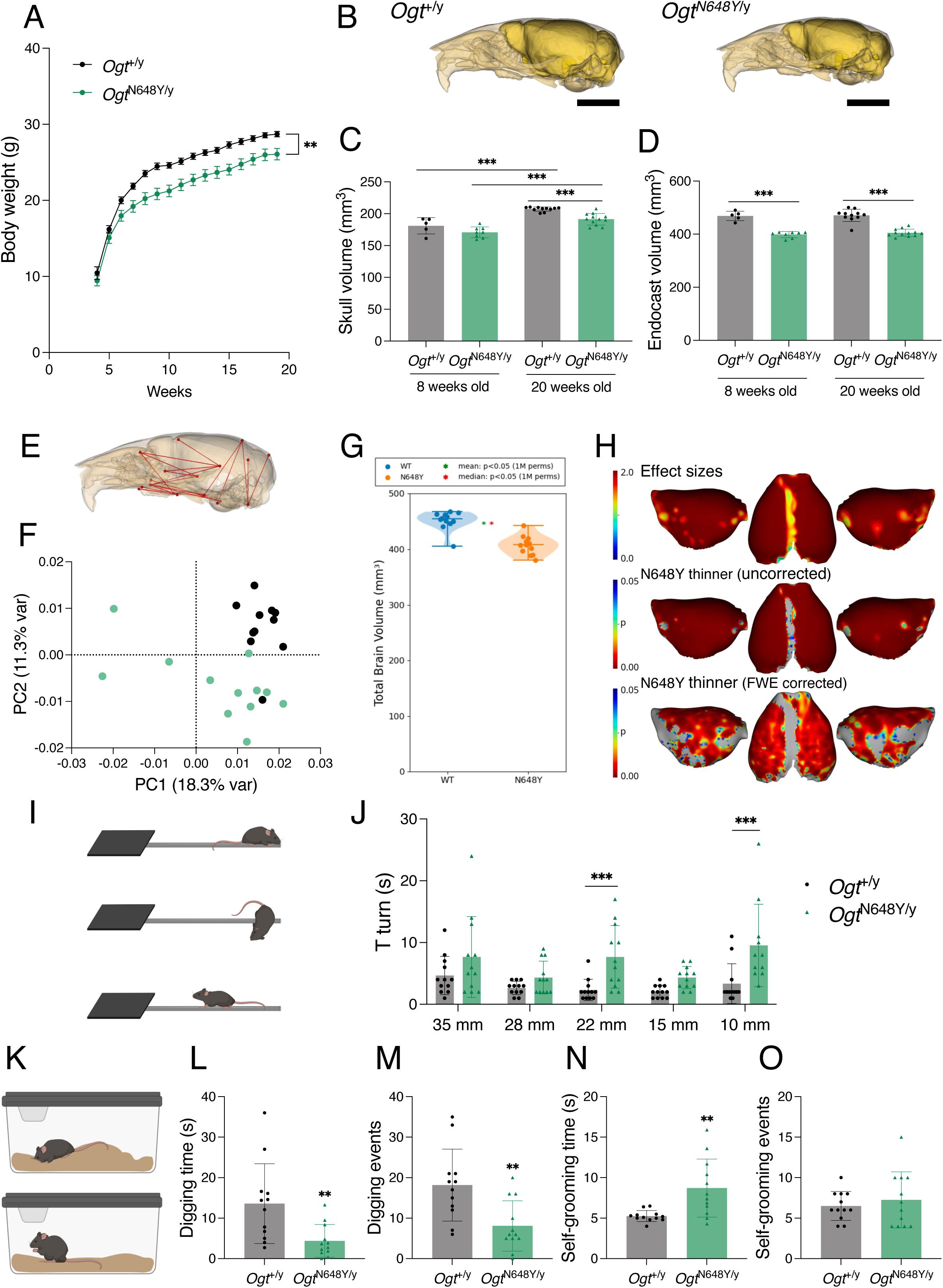
OGT^N648Y^ variant leads to microcephaly and motor deficits in mice. Data are represented as mean ± SD. The n-numbers are n = 5 and n = 8 for the 8-week-old *Ogt*^+/y^ and *Ogt*^N648Y/y^, respectively, and n = 12 for both genotypes at 20 weeks old. One *Ogt*^+/y^ skull was excluded from microCT analysis due to damage during sample processing. Student *t*-test is used for statistics unless another test is indicated. Significance shown as: \**p* < 0.05, \*\**p* < 0.01 and \*\*\**p* < 0.001 **(a)** Measurement of body weight over time of 12 *Ogt*^+/y^ and 12 *Ogt*^N648Y/y^ mice. Two-way ANOVA (alpha, 0.05) is used for statistics. **(b)** Lateral views of representative 3D reconstructions of the skull with endocast of *Ogt*^+/y^ (left) and *Ogt*^N648Y/y^ (right) mice. Scale bar: 5 mm. **(c)** Volume of the skulls of 8- and 20-week-old *Ogt*^+/y^ and *Ogt*^N648Y/y^ mice. **(d)** Volume of the endocasts of 8- and 20-week-old *Ogt*^+y^ and *Ogt*^N648Y/y^ mice. **(e)** Lateral view of skull indicating linear distances at least 5% shorter in *Ogt*^N648Y/y^ mice compared to *Ogt*^+/y^. **(f)** PCA biplot of PC1 and PC2 of the surface landmarks of 20-week-old *Ogt*^+/y^ (black) and *Ogt*^N648Y/y^ (light green) mice. Percentages in the axis indicate the explained variance by the corresponding PC. **(g)** Total brain volume obtained by MRI. Statistical significance from permutation tests is indicated by asterisks (*, green for group mean, red for median). **(h)** Statistical maps of group differences in cortical thickness between *Ogt*^+/y^ and *Ogt*^N648Y/y^. Maps are effect sizes (top row), and p-values of significant differences, both uncorrected for multiple comparisons (middle row), and with family-wise error (FWE) correction. **(i)** Pictogram illustrating the static rods test. Mice are placed at the end of the rod (top), and the time required to perform the T-turn (middle) and reach the platform (bottom) is recorded. Created with BioRender. **(j)** Time to perform the T-turn of *Ogt*^+/y^ and *Ogt*^N648Y/y^ during the static rod tests. **(k)** Pictogram illustrating the digging and self-grooming tests. Created with BioRender. **(l)** Time spent digging by *Ogt*^+/y^ and *Ogt*^N648Y/y^ during 3 min observation. **(m)** Number of digging events by *Ogt*^+/y^ and *Ogt*^N648Y/y^ during 3 min observation. **(n)** Time spent self-grooming by *Ogt*^+/y^ and *Ogt*^N648Y/y^ during 3 min observation. **(o)** Number of self-grooming events by *Ogt*^+/y^ and *Ogt*^N648Y/y^ during 3 min observation.

### Ogt^N648Y/y^ mice exhibit sensory-motor deficits

Patients with OGT-CDG variants experience cognitive and motor impairments (50). Therefore, we performed extensive behavioural screening of the *Ogt*^N648Y/y^ mice, including locomotion, memory, anxiety, and compulsive assessments to establish quantifiable phenotypic differences with their WT littermates. We first assessed general locomotion activity in an open field arena, where mice are placed in an open arena for 10 minutes for three consecutive days to evaluate exploratory drive in both novel and familiar environments. We observed an increase in the distance travelled and the maximum speed over the three days in *Ogt*^N648Y/y^ mice compared to their WT littermates (**Fig. S3A-C**).

We next assessed sensory motor and coordination skills in the *Ogt*^N648Y/y^ mice using static rod tests, where mice are placed at the end of horizontal rods of decreasing diameters and are required to perform a T-turn before reaching a safety platform (**Fig. 2I**). We observed that although all *Ogt*^N648Y/y^ mice were able to perform the test, they required more time compared to WT mice as seen by an increase in total time for the 28, 22, 15, and 10 mm diameter rods (**Fig. S3D).** This difference was driven by an increase in time to perform the motorically challenging T-turn in the *Ogt*^N648Y/y^ mice compared to WT (**Fig. 2J**), while the subsequent time to reach the safety platform was similar for both genotypes (**Fig. S3E**).

In addition, we assessed anxiety-like behaviour in the *Ogt*^N648Y/y^ mice using the elevated plus maze (EPM) test (61). We observed that *Ogt*^N648Y/y^ mice showed similar anxiety levels compared to WT mice, as seen by similar time spent in open and closed arms (**Fig. S3F-I**), and comparable time spent in the light compartment in the dark-light paradigm (**Fig. S3J-M**) between both genotypes. Of note, *Ogt*^N648Y/y^ travelled a greater distance and displayed more entries during the initial 5 min period of the dark-light test compared to WT. Similarly, *Ogt*^N648Y/y^ showed a mild increase in distance travelled and centre entries in the EPM test, albeit not significant.

We then assessed repetitive behaviour in the *Ogt*^N648Y/y^ mice. During a 3-minute observation period (**Fig. 2K**), we observed a decrease both in the time spent digging (**Fig. 2L**) and in the number of digging events (**Fig. 2M**) in the *Ogt*^N648Y/y^ mice compared to WT littermates. During that same period, we observed an increase in the time spent self-grooming in the *Ogt*^N648Y/y^ mice (**Fig. 2N**), while the number of self-grooming events did not differ from those of the WT (**Fig. 2O**). We did not see any difference between *Ogt*^N648Y/y^ and WT mice in the number of buried marbles (**Fig. S3N**). Additionally, during both the nesting and burrowing tube tests, *Ogt*^N648Y/y^ mice removed the same amount of material as WT (**Fig. S3O, P**).

Finally, we evaluated cognitive function in the mice by assessing novelty-associated memory using the novel object recognition (NOR) test. We did not observe a significant difference in the discrimination indexes for both short- and long-term paradigms (**Fig. S3Q, R**), suggesting intact novelty-associated memory in the *Ogt*^N648Y/y^ mice. Taken together, these observations suggest that *Ogt*^N648Y/y^ mice show an impairment in fine sensory motor skills, accompanied by altered exploratory behaviour and a degree of increased activity.

### Genetic rescue of O-GlcNAc dyshomeostasis in Ogt^N648Y/y^ mice

Both *Ogt*^N648Y/y^, described here, and *Ogt*^C921Y/y^ (61) OGT-CDG mouse models caused impaired brain O-GlcNAc dyshomeostasis. Lower O-GlcNAc levels and altered OGA/OGT ratios were observed, which could underpin the morphological and behaviour deficits observed in the mouse models, although it is not yet clear whether these deficits are of neurodevelopmental or neurophysiological origin. Targeting OGA activity using small molecules has been widely used in neurodegenerative diseases to raise O-GlcNAc levels and improve cognitive function in models of neurodegeneration (32,34,39). In an attempt to genetically validate this approach and rescue pathogenic O-GlcNAc dyshomeostasis in the *Ogt*^N648Y/y^ mice, we made use of the previously generated *Oga*^D285A^ mouse model (15). In the *Oga*^D285A^ mice, the key Asp285 residue that is required for both catalytic activity and O-GlcNAc binding has been replaced with Ala, leading to abolition of O-GlcNAcase activity and increased O-GlcNAc levels. Therefore, these mice can be exploited in a genetic strategy to model chronic inhibition of OGA activity from conception to adulthood. As homozygous *Oga*^D285A/D285A^ exhibit perinatal lethality, we used heterozygous *Oga*^D285A/+^ mice for subsequent breeding. Firstly, we assessed the viability and Mendelian inheritance distribution of male offspring generated from heterozygous *Ogt*^N648Y/+^ females crossed with heterozygous *Oga*^D285A/+^ males. DNA from the males in the offspring was genotyped and sequenced. Hemizygous *Ogt*^N648Y/y^ males, heterozygous *Oga*^D285A/+^ males, and the compound heterozygous *Ogt*^N648Y/y^*Oga*^D285A/+^ males could be obtained. We then analysed Mendelian inheritance of WT *Ogt*^+/y^*Oga*^+/+^, heterozygous *Ogt*^+/y^*Oga*^D285A/+^, hemizygous *Ogt*^N648Y/y^*Oga*^+/+^, and compound heterozygous *Ogt*^N648Y/y^*Oga*^D285A/+^ male animals obtained from the crosses. Compared to the expected 25% per genotype, we observed that the crosses generated 30 % *Ogt*^+/y^*Oga*^+/+^, 29 % *Ogt*^+/y^*Oga*^D285A/+^, 21 % *Ogt*^N648Y/y^*Oga*^+/+^, and 21 % *Ogt*^N648Y/y^*Oga*^D285A/+^ male pups (Chi-square = 2.3, n = 73) (**Fig. 3A**), suggesting absence of lethality of the double *Ogt*^N648Y/y^*Oga*^D285A/+^ mutant animals and inheritance patterns similar to the *Ogt*^N648Y/y^ alone (**Fig. 1C**).

**Figure 3:**
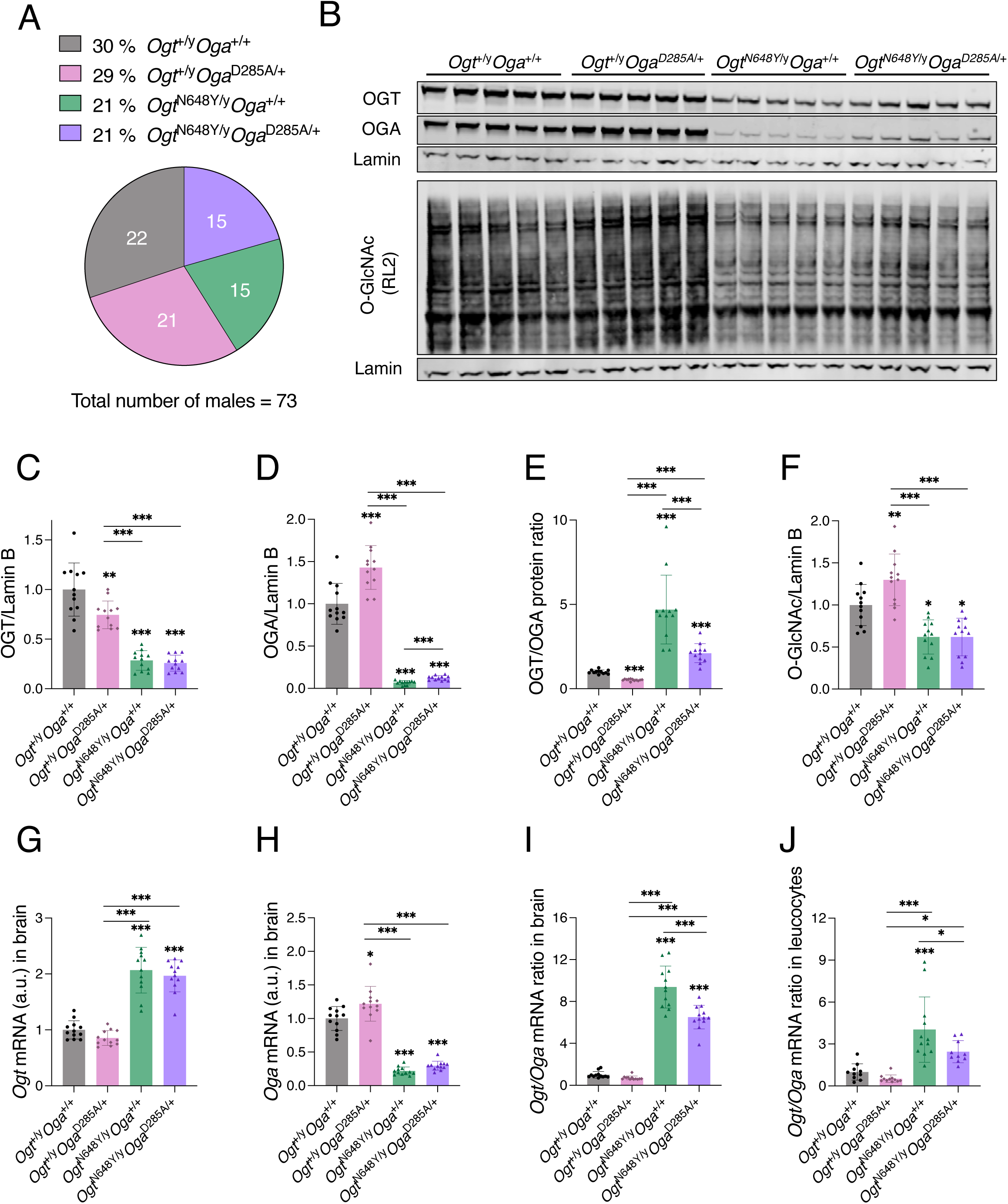
Genetic rescue of O-GlcNAc dyshomeostasis by crossing *Ogt*^N648Y/y^ mice with OGA-deficient mice. Data are represented as mean ± SD. The n-number is n = 12 for all genotypes. One-way ANOVA (alpha, 0.05) is used for statistics. Significance shown as: \**p* < 0.05, \*\**p* < 0.01 and \*\*\**p* < 0.001. **(a)** Diagram indicating percentages and numbers of male *Ogt*^+/y^*Oga*^+/+^, *Ogt*^+/y^*Oga*^D285A/+^, *Ogt*^N648Y/y^*Oga*^+/+^, and *Ogt*^N648Y/y^*Oga*^D285A/+^ animals generated from female *Ogt*^N648Y/+^*Oga*^+/+^ x male *Ogt*^+/y^*Oga*^D285A/+^ breeding pairs. **(b)** Western blot of OGT, OGA, and O-GlcNAcylated protein levels in adult brain of *Ogt*^+/y^*Oga*^+/+^, *Ogt*^+/y^*Oga*^D285A/+^, *Ogt*^N648Y/y^*Oga*^+/+^ and *Ogt*^N648Y/y^*Oga*^D285A/+^ mice. Lamin antibody was used as a loading control. **(c)** Quantification of OGT protein levels from the Western blot in panel (b). **(d)** Quantification of OGA protein levels from the Western blot in panel (b). **(e)** Quantification of OGT/OGA protein ratio from the Western blot in panel (b) **(f)** Quantification of O-GlcNAcylated proteins levels from the Western blot in panel (b). **(g)** Quantification of *Ogt* mRNA in whole mouse brain of *Ogt*^+/y^*Oga*^+/+^, *Ogt*^+/y^*Oga*^D285A/+^, *Ogt*^N648Y/y^*Oga*^+/+^ and *Ogt*^N648Y/y^*Oga*^D285A/+^ mice by RT-PCR. **(h)** Quantification of *Oga* mRNA in whole mouse brain of *Ogt*^+/y^*Oga*^+/+^, *Ogt*^+/y^*Oga*^D285A/+^, *Ogt*^N648Y/y^*Oga*^+/+^ and *Ogt*^N648Y/y^*Oga*^D285A/+^ mice by RT-PCR. **(i)** Quantification of *Ogt/Oga* mRNA ratio in whole mouse brain of *Ogt*^+/y^*Oga*^+/+^, *Ogt*^+/y^*Oga*^D285A/+^, *Ogt*^N648Y/y^*Oga*^+/+^ and *Ogt*^N648Y/y^*Oga*^D285A/+^ mice by RT-PCR. **(j)** Quantification of *Ogt/Oga* mRNA ratio in leucocytes of *Ogt*^+/y^*Oga*^+/+^, *Ogt*^+/y^*Oga*^D285A/+^, *Ogt*^N648Y/y^*Oga*^+/+^ and *Ogt*^N648Y/y^*Oga*^D285A/+^ mice by RT-PCR.

We then evaluated the morphological parameters of the *Ogt*^N648Y/y^*Oga*^D285A/+^ mice. Both *Ogt*^N648Y/y^ and *Ogt*^N648Y/y^*Oga*^D285A/+^ animals exhibited reduced body weight, body length, skull size, and brain weight compared to their WT and *Oga*^D285A/+^ littermates (**Fig. S4**). No differences between *Ogt*^N648Y/y^ and *Ogt*^N648Y/y^*Oga*^D285A/+^ genotypes were observed for all measured parameters, suggesting no restoration of body morphometrics in the *Ogt*^N648Y/y^*Oga*^D285A/+^ animals. To investigate whether *Ogt*^N648Y/y^*Oga*^D285A/+^ mice will show improved behaviour parameters, we performed static rods and digging tests, for which pronounced phenotypes were observed in *Ogt*^N648Y/y^ mice **(Fig. 2I-O).** We also included gait analysis to detect ataxia and gait abnormalities that have been reported in patients carrying both *Ogt*^N648Y/y^ and *Ogt*^C921Y/y^ variants (51,58). Similar T-turn times were observed between both *Ogt*^N648Y/y^ and *Ogt*^N648Y/y^*Oga*^D285A/+^ genotypes during the static rods test (**Fig. S5A, B**). Similarly, no differences in digging or self-grooming time and event activities were observed between *Ogt*^N648Y/y^ and *Ogt*^N648Y/y^*Oga*^D285A/+^ mice (**Fig. S5C-F**). Although both *Ogt*^N648Y/y^ and *Ogt*^N648Y/y^*Oga*^D285A/+^ mice exhibited shorter stride length, narrower step width, and impaired step alternation (**Fig. S5G-L**) compared to WT littermates in gait analysis, no differences were observed after normalisation with body length between the four genotypes for all parameters (**Fig. S5M-Q**), suggesting unaltered gait in both *Ogt*^N648Y/y^ and *Ogt*^N648Y/y^*Oga*^D285A/+^ mice.

We next examined the effect of genetic loss of OGA activity on O-GlcNAc homeostasis in the *Ogt*^N648Y/y^*Oga*^D285A/+^ mice. We assessed levels of OGT, OGA, and O-GlcNAc by western blotting (**Fig. 3B**). OGT protein levels in the double mutant remained mostly unchanged compared to *Ogt*^N648Y/y^ mice, however the amount of OGA protein increased (**Fig. 3C, D**). OGT/OGA protein ratio in *Ogt*^N648Y/y^*Oga*^D285A/+^ mice was lower compared to *Ogt*^N648Y/y^ mice (**Fig. 3E**), suggesting a partial rescue of imbalance of the O-GlcNAc cycling enzymes, although this did not affect overall O-GlcNAcylation (**Fig. 3F**). To further explore this, we measured *Ogt* and *Oga* mRNA expression in whole brains from *Ogt*^N648Y/y^*Oga*^D285A/+^ mice and their littermates, showing that *Ogt* mRNA expression is unchanged in *Ogt*^N648Y/y^*Oga*^D285A/+^ mice compared to *Ogt*^N648Y/y^, while *Oga* mRNA expression appears to be slightly increased (**Fig. 3G, H**). When combining both measurements, we found that the *Ogt/Oga* ratio in the brain was significantly lower in *Ogt*^N648Y/y^*Oga*^D285A/+^ mice compared to *Ogt*^N648Y/y^ littermates (**Fig. 3I**), similar to the observations at the protein level. We also analysed *Ogt* and *Oga* mRNA expression in the blood of the same mice as a proxy of brain O-GlcNAc dyshomeostasis. Indeed, the *Ogt/Oga* ratio was also lower in blood of *Ogt*^N648Y/y^*Oga*^D285A/+^ mice compared to *Ogt*^N648Y/y^ littermates (**Fig. 3J; S6A, B**). Overall, these findings suggest that although we did not observe rescue of morphological and behavioural deficits in the *Ogt*^N648Y/y^*Oga*^D285A/+^ mice, genetic inhibition of OGA did achieve partial rescue of brain O-GlcNAc dyshomeostasis that can be indirectly measured as a biomarker in peripheral blood.

## Discussion

Pathogenic variants in OGT have been linked to ID, resulting in the recently described OGT-CDG syndrome (50,56–58). Affected individuals exhibit a wide range of physical, neurological, and behavioural symptoms, with varying degrees of prevalence and severity. OGT-CDG variants are mostly *de novo* mutations found in families worldwide, contributing to the heterogeneity of the disorder. Animal models are valuable for studying disease aetiology, uncovering underlying mechanisms, and developing treatments. Therefore, to dissect the molecular basis of OGT-CDG pathology and establish a platform for evaluating possible treatments of the disease, vertebrate OGT-CDG models are required to establish patient-linked, reliably quantifiable phenotypes. Due to the heterogeneity in symptoms and the number of reported variants, multiple models are necessary to fully recapitulate the disease and expand the identification of phenotypes for targeted therapeutic interventions. Previously, we successfully generated and described a mouse carrying an OGT-CDG variant in the catalytic core, *Ogt*^C921Y/y^ (56,61), which shows altered O-GlcNAc homeostasis, reduced body and brain size, and behavioural abnormalities, including hyperactivity and impulsivity. In the present study, we provided a biochemical, morphological, and behavioural characterization of a newly generated mouse line with the N648Y variant in the catalytic core of OGT and used a genetic approach to correct O-GlcNAc dyshomeostasis.

The OGT^N648Y^ variant is a *de novo* variant in *OGT* found in the second child of an Estonian non-consanguineous couple (51). The patient exhibited a range of symptoms similar to those found in other OGT-CDG patients, such as intellectual disability, dysmorphic features, and developmental delay (50). *Ogt*^N648Y/y^ mice exhibit characteristic morphological features such as lower body weight and size, also often observed in other mouse models of X-linked IDs, where the mice show decreased body weight but remain healthy and fertile (62). *Ogt*^N648Y/y^ mouse skulls are smaller in size and differ in shape compared to WT. A number of OGT-CDG patients show skull asymmetry and microcephaly (50,51), clinically described as a reduction in the head circumference, which suggests a decrease in brain volume (63). *Ogt*^N648Y/y^ mice display a microcephaly phenotype that is stronger than that observed in the other OGT-CDG line currently available (56,61), and the brains of *Ogt*^N648Y/y^ mice are smaller than those of WT mice. However, as opposed to *Ogt*^C921Y/y^ mice, all brain regions in *Ogt*^N648Y/y^ mice are globally reduced, with no regions differentially smaller after normalizations. *Ogt*^N648Y/y^ mice behave differently from WT mice in the static rods test, particularly regarding the performance of the T-turn, which suggests defects in motor coordination and balance. Furthermore, they show reduced digging activity, while other tests assessing compulsive behaviour are not affected. The previously described *Ogt*^C921Y/y^ mice exhibited similar impairments in the static rod tests and digging activity (61). In other mouse models of IDs and autism spectrum disorders, a decrease in the digging activities was also observed, together with an increase in self-grooming (64–67). The decrease in free digging may indicate a lower anxiety level, which would align with the increase in transitions to the light compartment in the dark/light test, but can also be a measurement of lower exploratory drive (68). Therefore, *Ogt*^N648Y/y^ mice exhibit motor defects and autistic spectrum disorder-related behaviours, features that recapitulate patients’ characteristics (57). Notably, during OF, EPM, and dark/light tests, the *Ogt*^N648Y/y^ mice show a tendency to increase the distance travelled, which may indicate mild hyperactivity, a characteristic also reported for the patient carrying the OGT^N648Y^ variant and more evident in the previously described *Ogt*^C921Y/y^ mice (61). Therefore, traits of the patients appear to be partially reflected in this mouse model. It remains to be seen whether different variants in the OGT gene causing ID may impact mice differently. Taken together, this work establishes quantifiable morphometric and behavioural phenotypes in an OGT-CDG mouse model that can be used to dissect mechanisms and evaluate future treatments.

To date, several mechanisms have been proposed to explain the disease, including disruptions in O-GlcNAc homeostasis, HCF1 processing, or misfolding of the OGT protein (50). *Ogt*^N648Y/y^ mice exhibit reduced global O-GlcNAc levels in the brain, in agreement with what was observed in mouse embryonic stem cells carrying the mutation (51) and in other models carrying the OGT^C921Y^ variant (56,58,59). Both OGT and OGA protein levels appear to be lower in the brains of the *Ogt*^N648Y/y^ mouse, but the reduction in OGT protein levels is not explained by reduced transcription, as *Ogt* mRNA levels are, in fact, increased, presumably as a compensatory mechanism. X-ray crystallography performed with the OGT^N648Y^ variant indicates structural rearrangements in the catalytic domain, suggesting that OGT protein folding or stability may be affected (51). These observations, along with the reduction in *Oga* mRNA levels, were likewise observed in other OGT-CDG mice (35) and suggest a compensatory mechanism to counteract the pathogenic hypoglycosylation. Taken together, these findings suggest O-GlcNAc dyshomeostasis as a common hallmark in OGT-CDG mice.

Several neurological diseases are linked to disrupted O-GlcNAc balance, and there are multiple cases where OGA inhibition has been suggested as a possible treatment approach (32,34,39,41,42,46). Here, we use a genetic approach to restore O-GlcNAc levels in *Ogt*^N648Y/y^ mice using *Oga*^D285A/+^ mice, aiming to mimic chronic OGA inhibition from embryogenesis to adulthood in the context of OGT-CDG. *Oga*^D285A/+^ mice exhibit increased OGA levels and reduced OGT protein expression, likely as a response to the deficiency in OGA catalytic activity (15). Meanwhile, total O-GlcNAc levels remain elevated due to the reduction of OGA activity. As homozygous *Oga*^D285A/D285A^ mice do not survive after birth, we used heterozygous *Oga*^D285A/+^ mice for modulating O-GlcNAc homeostasis in *Ogt*^N648Y/y^ mice. Heterozygous *Oga*^D285A/+^ mice develop normally (15), and in agreement with this, we show here that heterozygous *Oga*^D285A/+^ mice display similar morphometric parameters and behaviour to wild type mice in all tests conducted, suggesting that moderate, chronic, longitudinal elevation of O-GlcNAcylation levels is in principle a safe approach for OGT-CDG phenotypic rescue. Moreover, an increase in O-GlcNAc levels has been shown not to affect the structure and function of neurons and to provide a neuroprotective effect against diseased or aging brains (33,37,69). In *Ogt*^N648Y/y^*Oga*^D285A/+^ mouse brains, we achieved a partial rescue of OGT/OGA homeostasis, driven by an increase in OGA protein levels and mRNA expression in response to heterozygotic loss of OGA activity. However, the partial correction of the OGT/OGA ratio was not sufficient to restore O-GlcNAc levels, behavioural deficits, and morphometrics in these mice. The use of heterozygous mice due to homozygotic lethality constitutes one limitation of our approach, as heterozygous mice still produce a functional copy of OGA, which may compensate for the heterozygotic loss of OGA function. Previous studies indicated that it is necessary to inhibit OGA by more than 80% to observe neuroprotective hyper-O-GlcNAcylation in the brains of mouse models of neurodegenerative diseases (32,34,70). Although our genetic approach did not rescue OGT-CDG phenotypes, it demonstrates the validity of OGA targeting in modulating OGT/OGA homeostasis in OGT-CDG from conception without leading to deleterious effects. Future avenues for modulating O-

GlcNAc homeostasis in OGT-CDG mice may include the use of small molecules to achieve greater OGA inhibition and metabolic approaches. Recently, nutrient supplementation in different CDG models has been studied as a treatment option (71–74), suggesting that using sugars that increase protein O-GlcNAcylation may be beneficial in the context of OGT-CDG. Similarly, supplementation with GlcNAc itself has recently been shown to have beneficial effects in multiple sclerosis patients in a small clinical trial (75). Another metabolite of interest is glucosamine, which can enter the hexosamine biosynthetic pathway, leading to increased UDP-GlcNAc levels and, consequently, directly boost protein O-GlcNAc levels (76,77). Glucosamine is a widely used supplement for joint health (78,79), and it has also been suggested to be beneficial in several metabolic diseases (80,81). Supplementation with glucosamine has already been explored in the patient carrying the OGT^N648Y^ variant and may have a positive impact without any adverse effects (51). OGT-CDG mice may serve as a crucial model to better understand the impact of metabolite supplementation in the context of this disease.

Biomarkers serve as measurable and accessible indicators of both disease traits and the impact of a treatment or intervention. In OGT-CDG, such biomarkers are urgently needed to evaluate the effect of genetic or pharmacological rescue strategies. Recently, O-GlcNAc dyshomeostasis has been proposed as a biomarker for identifying new pathogenic OGT-CDG variants using a stem cell reporter line (82,83). Importantly, peripheral blood has been used to identify O-GlcNAc-related changes in both disease and treatment contexts. In a study involving an OGA inhibitor as treatment for proteinopathies, changes in protein O-GlcNAcylation were detected in both brain and blood (84). Similarly, in patients with diabetes mellitus, OGA expression and activity in leucocytes correlated with systemic inflammation (85). Here, we observed a partial rescue of *Ogt/Oga* mRNA ratio in the brain that was also detectable in leucocytes extracted from peripheral blood. These findings highlight the potential of using blood *Ogt*/*Oga* mRNA ratio not only for OGT-CDG pathogenicity detection but also as a biomarker to evaluate treatment-induced modulation of O-GlcNAc dyshomeostasis in OGT-CDG disorder.

In conclusion, the *Ogt*^N648Y/y^ mouse line recapitulates key features of OGT-CDG, including disrupted O-GlcNAc homeostasis, growth deficits, and behavioural changes, making it a valuable model for dissecting disease mechanisms. Although the genetic rescue approach used here only partially restored O-GlcNAc balance in the brain without improving behaviour or morphology, the molecular changes were also observed in peripheral blood, supporting its use as an accessible biomarker for the disease and treatment response. Future research will focus on exploiting these OGT-CDG mouse models and biomarkers to evaluate therapeutic strategies for this patient population with unmet medical need.

## Materials & Methods

### Generation of mouse lines and animal husbandry

*Ogt*^N648Y/y^ mice were produced by microinjection as described previously (42). Genomic DNA from offspring was genotyped and sequenced to confirm the presence of the OGT^N648Y^ variant. Editing CRISPR reagents, Cas9 Nickase (1081062; Integrated DNA Technologies), and primers were purchased from Integrated DNA Technologies, and sequences are listed in **Table S1**. Founder *Ogt*^N648Y/y^ mice were crossed to C57BL/6JRj WT animals (Janvier, France) for further breeding. *Oga*^D285A^ mice were previously reported (15) and were maintained under C57BL/6JRj background. Mice carrying the double mutation (*Ogt*^N648Y/y^*Oga*^D285A/+^) were generated by crossing *Ogt*^N648Y/+^ females with heterozygous *Oga*^D285A/+^ males, leading to *Ogt*^N648Y/y^*Oga*^D285A/+^ animals. Mice were housed in ventilated cages with water and food available *ad libitum* and 12/12 h light/dark cycles in the Skou animal facility (Aarhus University). All animal studies and breeding were performed following the ARRIVE guidelines and the European Communities Council Directive (2010/EU) and were approved by the Danish Animal Experiments Inspectorate (Dyreforsøgstilsynet), under Breeding license: 2022-15-0202-00135 and Project license: 2023-15-0201-01426.

### Tissue collection and disruption for protein and RNA analysis

Mice were euthanised by intraperitoneal injection of pentobarbital (overdose). Brain tissue from male mice was rapidly dissected, snap frozen in liquid nitrogen, and stored at −80 °C. Tissues were dissociated in Phosphate-Buffered Saline (PBS) using Precellys^®^ 24 Touch homogenizer (Bertin Technologies) as described previously (61). Brain homogenates were split in half for further protein and RNA extractions.

### Leucocyte extraction from peripheral blood

Whole blood was collected in heparinized tubes from intracardiac puncture. 0.5 mL of blood was lysed in 1 mL of Erythrocyte Lysis Buffer (ELB) (155 mM NH4Cl, 10mM NaHCO3, 1mM EDTA, pH 7.4). After a short mix, samples were incubated on ice for 10 min and spun at 1500 g for 5 min at 4 °C. Pellets were resuspended in ELB, mixed, and spun at 1500 g for 5 min. Finally, pellets were resuspended in 350 μL lysis buffer from the RNeasy Micro kit (ref, Qiagen) and stored at -80 °C until further RNA processing.

### Western blot

Brain homogenates were lysed using 10x RIPA buffer (Cell Signaling), sonicated (10 s on/10 s off at 20% amplitude, 6 pulses), and centrifuged at 14,000 rpm for 30 min at 4 °C. Proteins were quantified, separated, and transferred to nitrocellulose membranes as described previously (61). The antibodies used were: anti-OGA (MGEA5) (1:500 dilution; HPA036141; Sigma), anti-O-GlcNAc (RL2) (1:500 dilution; NB300-524, Novus Biologicals), anti-OGT (F-12) (1:1000 dilution; sc-74546; Santa Cruz), mouse anti-Lamin B (1:10000 dilution; 66095-Ig; Proteintech), and rabbit anti-Lamin B (1:10000; 12987-1-AP; Proteintech). Blots were imaged using a Li-Cor Odyssey infrared imaging system (LI-COR), and signals were quantified using Empiria Studio software from the same manufacturer. Data were normalized to the mean of each WT replicate group and expressed as fold change relative to WT.

### RT-qPCR analysis

Total RNA was purified from brain homogenates, followed by reverse transcription and quantitative PCR as described previously (56,61). Samples were analysed in biological replicates with technical triplicates using the comparative Ct method. The threshold-crossing value was normalized to internal control transcripts (*18S*, *Actb*, and *Pgk1*). Primers used for qPCR analysis are listed in **Table S2**. Data were normalized to the mean of each WT replicate group and expressed as fold change relative to WT.

### Behavioural studies

All animals were handled daily by the experimenter for a week before behavior testing. Handling occurred in the experimental room with ambient lighting of 25 to 30 Lux. Animals were housed in groups of 2 to 3 individuals. Behavioral testing was conducted over five consecutive weeks, with the mice aged between 10 and 14 weeks. Animals were placed in the experimental room at least 30 minutes before testing began. Data analysis was performed blindly regarding to genotype. The sample size was determined based on previous experience and confirmed through Post-hoc Power Calculation using the ClinCalc online tool (https://clincalc.com/stats/Power.aspx). Behaviour tests to assess locomotion and motor coordination (open field, rotarod, static rods,), cognitive performance (novel object recognition), anxiety (elevated maze, dark/light test) and compulsive behaviour (marbles, digging and self-grooming, nesting) were performed as described previously (61). Gait analysis was performed using a 40 cm long, 5.5 cm wide runway with a dark box at the end, covered with a strip of paper. The day before the test, the mice were trained to walk through the illuminated runway towards the dark box in a straight line. On the test day, the underside of the forepaws (red) and hind paws (blue) were painted, allowing them to walk along the corridor afterwards. Several measurements between the footprints were analysed, including stride length, step width, rear paws and front paws base of support (RBOS and FBOS, respectively), and step alternation.

### Brain perfusion and sample preparation for structural analysis

Following behaviour studies, *Ogt*^N648Y/y^ mice (WT n = 12; *Ogt*^N648Y/y^ n = 12) were anesthetized and perfused fixed for MRI as described previously (61). Briefly, transcardial perfusion fixation was followed by decapitation, after which the mandible and extracranial tissue were removed from the skull to prevent imaging artifacts. Hereafter, the in-skull brains were stored in 4% paraformaldehyde solution for at least one week. Before imaging, the samples were washed in PBS for a minimum of 24 h to increase MRI signal by removing excess fixative (86).

### Magnetic resonance imaging (MRI) and image analysis

MRI scans were acquired on a 9.4 T preclinical system (BioSpec 94/20, Bruker Biospin, Ettlingen, Germany) with a bore-mounted 25 mm quadrature transmit-receive coil. Structural data were collected as described previously (61) and aligned using precise multi-atlas segmentation (87). For all scan types, data quality was confirmed through visual inspection, and samples were rescanned if necessary to ensure consistently high data quality for subsequent analyses. Volumetric analyses were performed as previously described (61) using an in-house pipeline. Briefly, high-resolution FLASH images were pre-processed with B1 inhomogeneity correction, denoising, intensity normalization, and spatial alignment to a C57BL/6J mouse template via linear and non-linear registration. Neuroanatomical labels were then transformed back to native space to extract absolute and relative regional volumes (region size as % of total brain volume), accounting for differences in brain size. Cortical thickness was measured as previously described (61), generating statistical maps of group differences by fitting a general model at each surface vertex (SurfStat, http://www.math.mcgill.ca/keith/surfstat/). Due to multiple comparisons, statistical maps were family-wise error (FWE) corrected using random field theory (88) with α = 0.001 as the cluster-defining threshold. All statistical maps were thresholded at *p* = 0.05, both uncorrected and corrected.

### Micro Computed Tomography (microCT)

Following MRI, the in-skull brain samples were imaged using µCT (Scanco Viva CT 80) as described previously in detail (61). Skulls were imaged and analysed in 3D Slicer (https://www.slicer.org), where skulls were isolated and exported as 3D models. Image processing was performed with the investigator blinded to the group distribution. For 3D visualization of differences between WT and *Ogt*^N648Y/y^ mice, models for the average shapes of WT and *Ogt*^N648Y/y^ mouse skulls were generated from the General Procrustes Analysis (GPA) module. Skull shape and size differences were assessed both by computing the bone volume and by measuring distances between the 45 surface landmarks and performing Euclidean Distance Matrix Analysis. One WT skull was excluded from analysis due to accidental damage during sample preparation.

### Statistical analysis

Unless otherwise noted, statistical analyses were performed with Prism 9. D’Agostino & Pearson, Shapiro–Wilk, and Kolmogorov-Smirnov normality tests were conducted to verify normality. For data meeting normality requirements, the unpaired *t*-test was used for pairwise comparisons of wild-type and mutant mouse data, and two-way ANOVA was used for multiple comparisons. For data sets that did not meet normality, the Mann-Whitney test was used for pairwise.

## Acknowledgements

This work was funded by a Wellcome Trust Investigator Award (110061), a Novo Nordisk Fonden Laureate award (NNF21OC0065969) and a Villum Fonden Investigator (00054496) to D.M.F.v.A. The Novo Nordisk Foundation is gratefully acknowledged for funding the Scanco µCT equipment as a part of the Aarhus X-ray Imaging Alliance (AXIA). We also thank Asad Jan for support in gait analysis.

## Author contributions

F.A. and D.M.F.v.A conceived the study; F.A., I.E.A, K.S.C., C.S.S, S.F.E, N.O. and J.S.T. performed experiments; A.T.F. performed molecular biology; F.A., I.E.A., K.S.C. and D.M.F.v.A. analysed data and F.A., I.E.A. and D.M.F.v.A. interpreted the data and wrote the manuscript with input from all authors.

## Supplementary Materials

**Figure S1:**
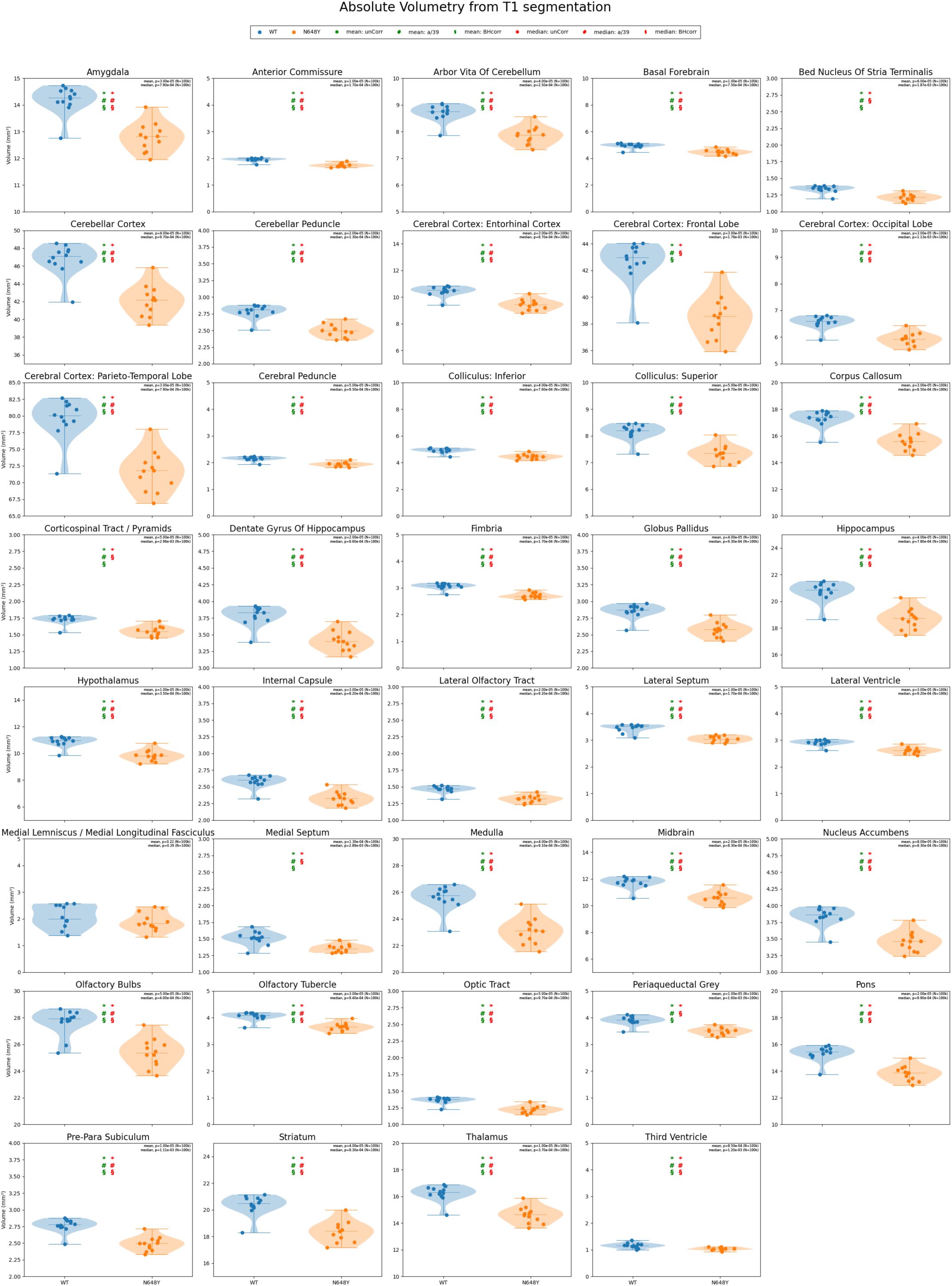
Absolute volumetry of 39 brain regions from T1. Each panel corresponds to a different bilaterally pooled region. Statistical significance (*p* < 0.05) from permutation tests (100k for each region) is indicated by asterisks (*; green for group mean, red for median) for uncorrected p-values. Correction for multiple comparisons was performed by division by the number of regions and is indicated by a pound symbol (#) or adjusted for false discovery rate (Benjamini-Hochsberg, BHcorr), indicated by a section sign (§) - if corrected, *p* < 0.05

**Figure S2:**
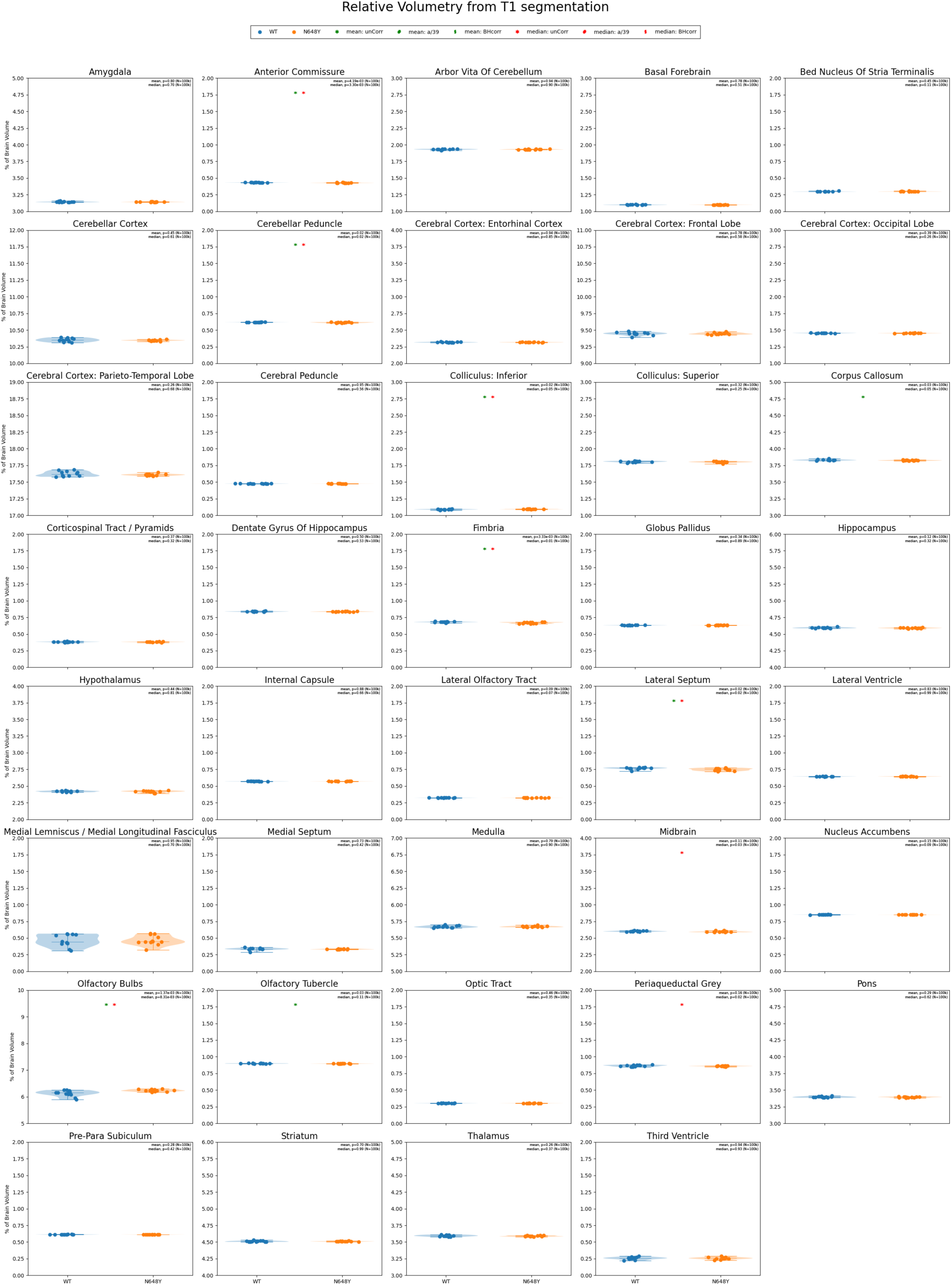
Relative volumetry of 39 brain regions from T1. Each panel corresponds to a different bilaterally pooled region. The regional relative volume (RRV) is normalized to the individual total brain volume. Each dot corresponds to a subject. Statistical significance (*p* < 0.05) from permutation tests (100k for each region) is indicated by asterisks (*; green for group mean, red for median) for uncorrected p-values. Correction for multiple comparisons was performed by division by the number of regions and is indicated by a pound symbol (#) or adjusted for false discovery rate (Benjamini-Hochsberg, BHcorr) indicated by a section sign (§) - if corrected *p* < 0.05.

**Figure S3:**
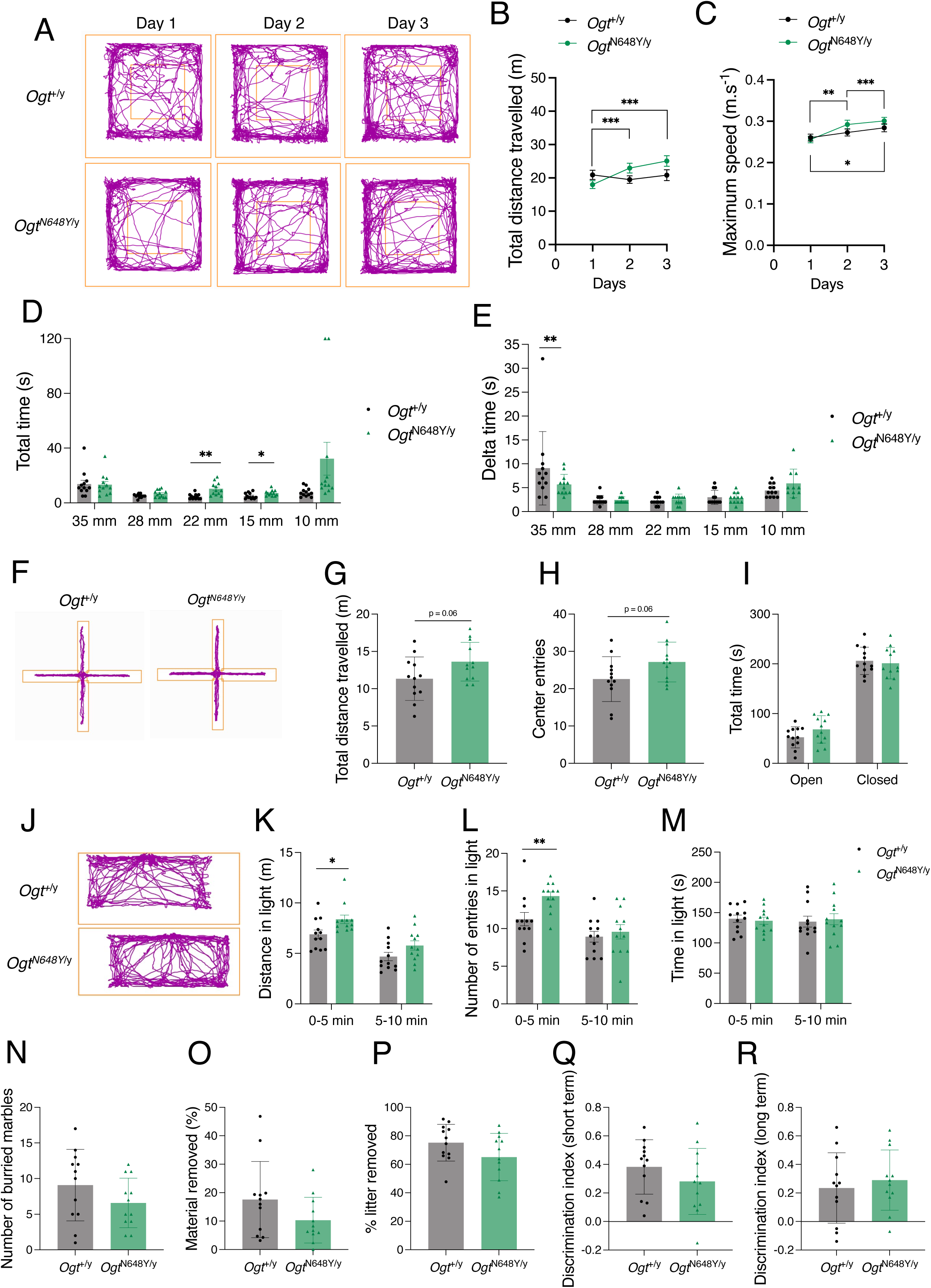
Behaviour of *Ogt*^N648Y/y^ mice. Data are represented as mean ± SD, n = 12 for all genotypes. Student’s t-test is used for statistics unless another test is indicated. Significance shown as: \**p* < 0.05, \*\**p* < 0.01 and \*\*\**p* < 0.001 **(a)** Representative tracking plot of one *Ogt*^+/y^ and one *Ogt*^N648Y/y^ mouse over three days in the open field arena **(b)** Total distance travelled over three consecutive days in the open field arena of *Ogt*^+/y^ (black) and *Ogt*^N648Y/y^ (green) mice. Two-way ANOVA (alpha, 0.05) was used for statistics. **(c)** Maximum speed achieved over three consecutive days in the open field arena by *Ogt*^+/y^ (black) and *Ogt*^N648Y/y^ (green) mice. Two-way ANOVA (alpha, 0.05) was used for statistics. **(d)** Total time to perform the static rods tests of *Ogt*^+/y^ (black) and *Ogt*^N648Y/y^ (green) mice. **(e)** Delta time (total time – time to perform the T-turn) of *Ogt*^+/y^ (black) and *Ogt*^N648Y/y^ (green) mice during the static rods test. **(f)** Representative tracking plot of one *Ogt*^+/y^ and one *Ogt*^N648Y/y^ mouse during the elevated plus maze (EPM) test. **(g)** Total distance travelled of *Ogt*^+/y^ and *Ogt*^N648Y/y^ mice during the EPM test. **(h)** Number of entries to the centre of *Ogt*^+/y^ and *Ogt*^N648Y/y^ mice during the EPM test. **(i)** Total time spent in open and closed arms of *Ogt*^+/y^ (black) and *Ogt*^N648Y/y^ (green) mice during the EPM test. **(j)** Representative tracking plot in the light compartment of one *Ogt*^+/y^ and one *Ogt*^N648Y/y^ mouse during the dark/light paradigm. **(k)** Distance travelled in the light of *Ogt*^+/y^ (black) and *Ogt*^N648Y/y^ (green) mice during the dark/light test. **(l)** Number of entries in the light of *Ogt*^+/y^ (black) and *Ogt*^N648Y/y^ (green) mice during the dark/light paradigm. **(m)** Time spent in the light of *Ogt*^+/y^ (black) and *Ogt*^N648Y/y^ (green) mice during the dark/light paradigm. **(n)** Number of buried marbles by *Ogt*^+/y^ and *Ogt*^N648Y/y^ mice during the marbles test. **(o)** Percentage of material removed by *Ogt*^+/y^ and *Ogt*^N648Y/y^ mice during the nesting test. **(p)** Percentage of litter removed by *Ogt*^+/y^ and *Ogt*^N648Y/y^ mice during the burrowing tube test. **(q)** Discrimination index of *Ogt*^+/y^ and *Ogt*^N648Y/y^ mice during the short-term (90 min) novel object (NOR) test. **(r)** Discrimination index of *Ogt*^+/y^ and *Ogt*^N648Y/y^ mice during the long-term (24 h) NOR test.

**Figure S4:**
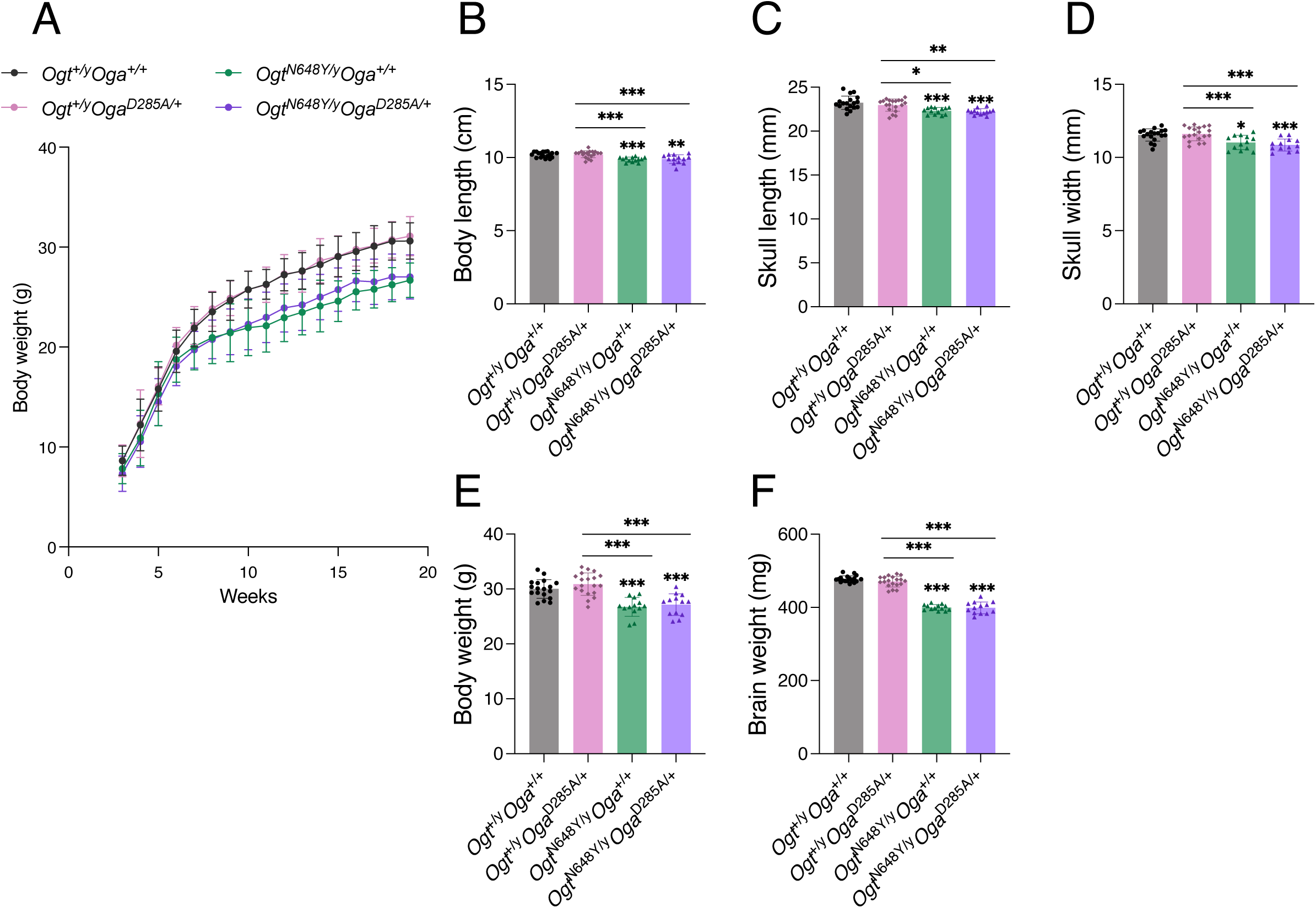
*Ogt*^N648Y/y^*Oga*^D285A/+^ mice morphometrics. Data for 20-week-old *Ogt*^+/y^*Oga*^+/+^, *Ogt*^+/y^*Oga*^D285A/+^, *Ogt*^N648Y/y^*Oga*^+/+^ and *Ogt*^N648Y/y^*Oga*^D285A/+^ mice unless otherwise stated. Data are represented as mean ± SD, n = 12 for all genotypes. One-way ANOVA (alpha, 0.05) is used for statistics unless another test is indicated. Significance shown as: \**p* < 0.05, \*\**p* < 0.01 and \*\*\**p* < 0.001. **(a)** Measurement of body weight over time. Two-way ANOVA (alpha, 0.05) is used for statistics. **(b)** Measurement of body length at 20 weeks. **(c)** Measurement of skull length at 20 weeks. **(d)** Measurement of skull width at 20 weeks. **(e)** Measurement of body weight at 20 weeks **(f)** Measurement of brain weight at 20 weeks.

**Fig. S5:**
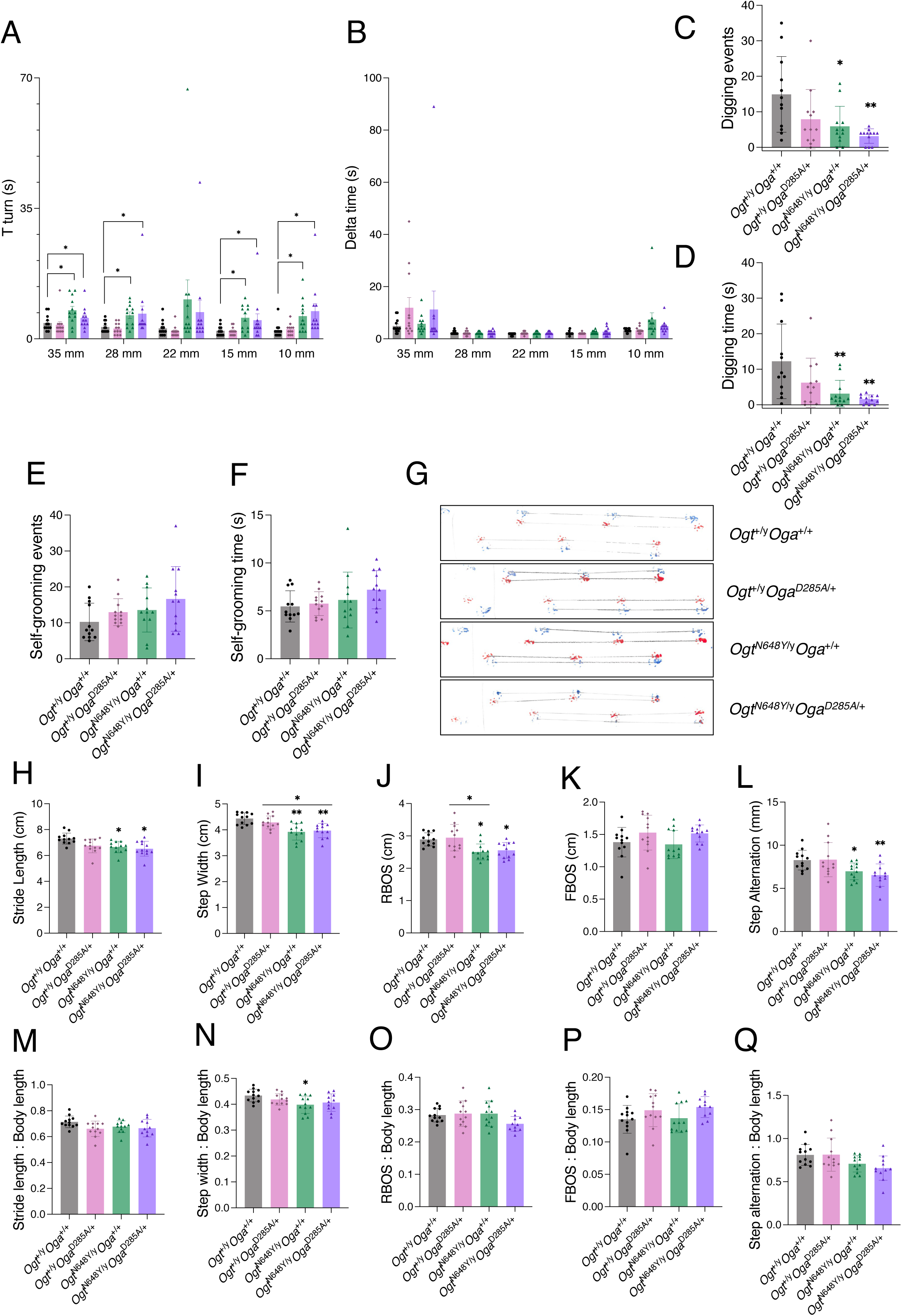
*Ogt*^N648Y/y^*Oga*^D285A/+^ mice behaviour tests. Data for behaviour tests performed in *Ogt*^+/y^*Oga*^+/+^, *Ogt*^+/y^*Oga*^D285A/+^, *Ogt*^N648Y/y^*Oga*^+/+^ and *Ogt*^N648Y/y^*Oga*^D285A/+^ mice. Data are represented as mean ± SD, n = 12 for all genotypes. One-way ANOVA (alpha, 0.05) is used for statistics. Significance shown as: \**p* < 0.05, \*\**p* < 0.01 and \*\*\**p* < 0.001. **(a)** Time to perform the T-turn of *Ogt*^+/y^*Oga*^+/+^ (black), *Ogt*^+/y^*Oga*^D285A/+^(pink), *Ogt*^N648Y/y^*Oga*^+/+^ (green) and *Ogt*^N648Y/y^*Oga*^D285A/+^ (purple) mice during the static rods test. **(b)** Delta time (total time – time to perform the T-turn) of *Ogt*^+/y^*Oga*^+/+^ (black), *Ogt*^+/y^*Oga*^D285A/+^(pink), *Ogt*^N648Y/y^*Oga*^+/+^ (green) and *Ogt*^N648Y/y^*Oga*^D285A/+^ (purple) mice during the static rods test. **(c)** Number of digging events during 3 min observation. **(d)** Time spent digging during 3 min observation. **(e)** Number of self-grooming events during 3 min observation. **(f)** Time spent self-grooming during 3 min observation. **(g)** Representative footprints of *Ogt*^+/y^*Oga*^+/+^, *Ogt*^+/y^*Oga*^D285A/+^, *Ogt*^N648Y/y^*Oga*^+/+^ and *Ogt*^N648Y/y^*Oga*^D285A/+^ mice. Red footprints correspond to the forepaws; blue footprints correspond to the hind paws. Lines represent measured distances. **(h)** Measurement of stride length as the distance between footprints of the same hind paw. **(i)** Measurement of step width as the contralateral distance between hind and front paws. **(j)** Measurement of rear paws base of support (RBOS) as the contralateral distance between hind paws. **(k)** Measurement of front paws base of support (FBOS) as the contralateral distance between forepaws. **(l)** Measurement of step alternation as the ipsilateral distance between hind and front paws. **(m)** Stride length normalised to body length. **(n)** Step width normalised to body length. **(o)** RBOS normalised to body length. **(p)** FBOS normalised to body length. **(q)** Step alternation normalised to body length.

**Figure S6:**
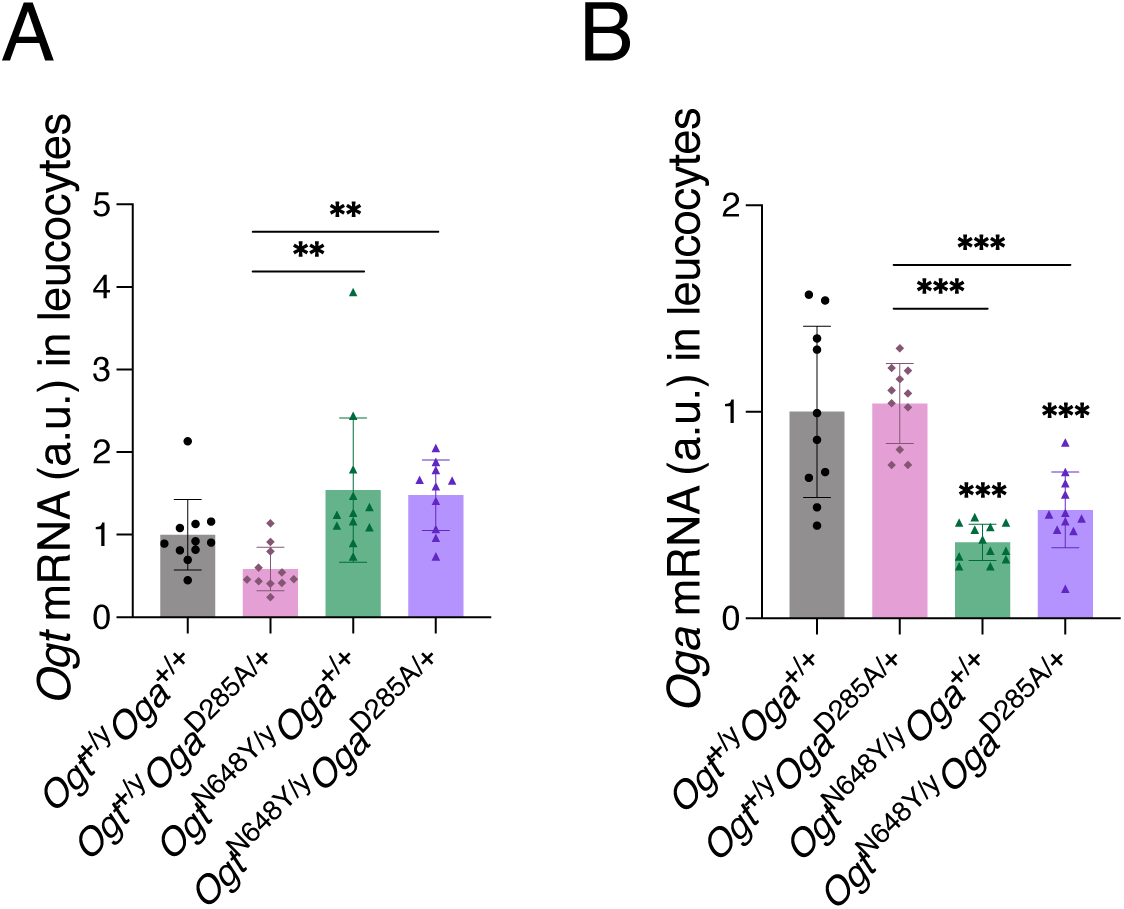
*Ogt*^N648Y/y^*Oga*^D285A/+^ blood biochemistry. Data for 20-week-old *Ogt*^+/y^*Oga*^+/+^ (n = 11), *Ogt*^+/y^*Oga*^D285A/+^ (n = 11), *Ogt*^N648Y/y^*Oga*^+/+^ (n = 12), and *Ogt*^N648Y/y^*Oga*^D285A/+^ (n = 11) mice. Data are represented as mean ± SD. One-way ANOVA (alpha, 0.05) is used for statistics. Significance shown as: \**p* < 0.05, \*\**p* < 0.01 and \*\*\**p* < 0.001 **(a)** Quantification of *Ogt* mRNA levels in blood by RT-PCR. **(b)** Quantification of *Oga* mRNA levels in blood by RT-PCR. One *Ogt*^+/y^*Oga*^+/+^ outlier was excluded from the analysis.

**Table S1:**
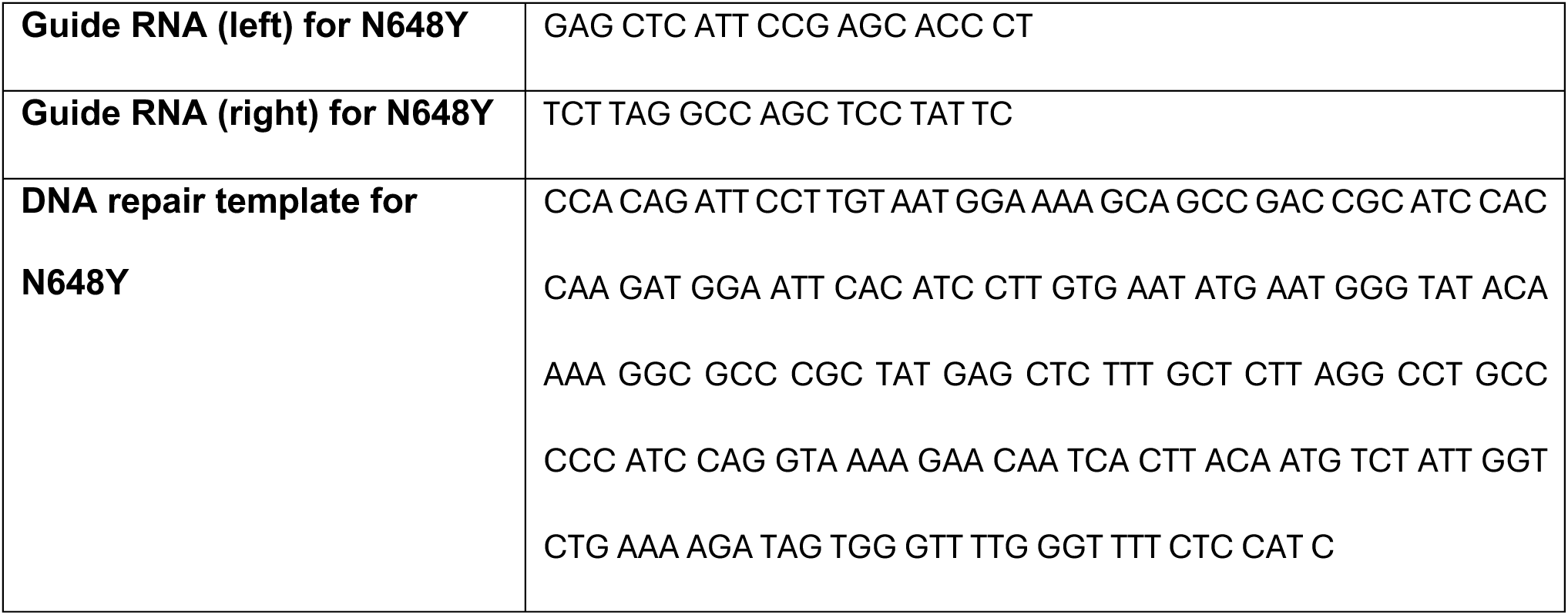
Sequences of reagents used for introducing the N648Y variant to *Ogt*.

**Table S2:**
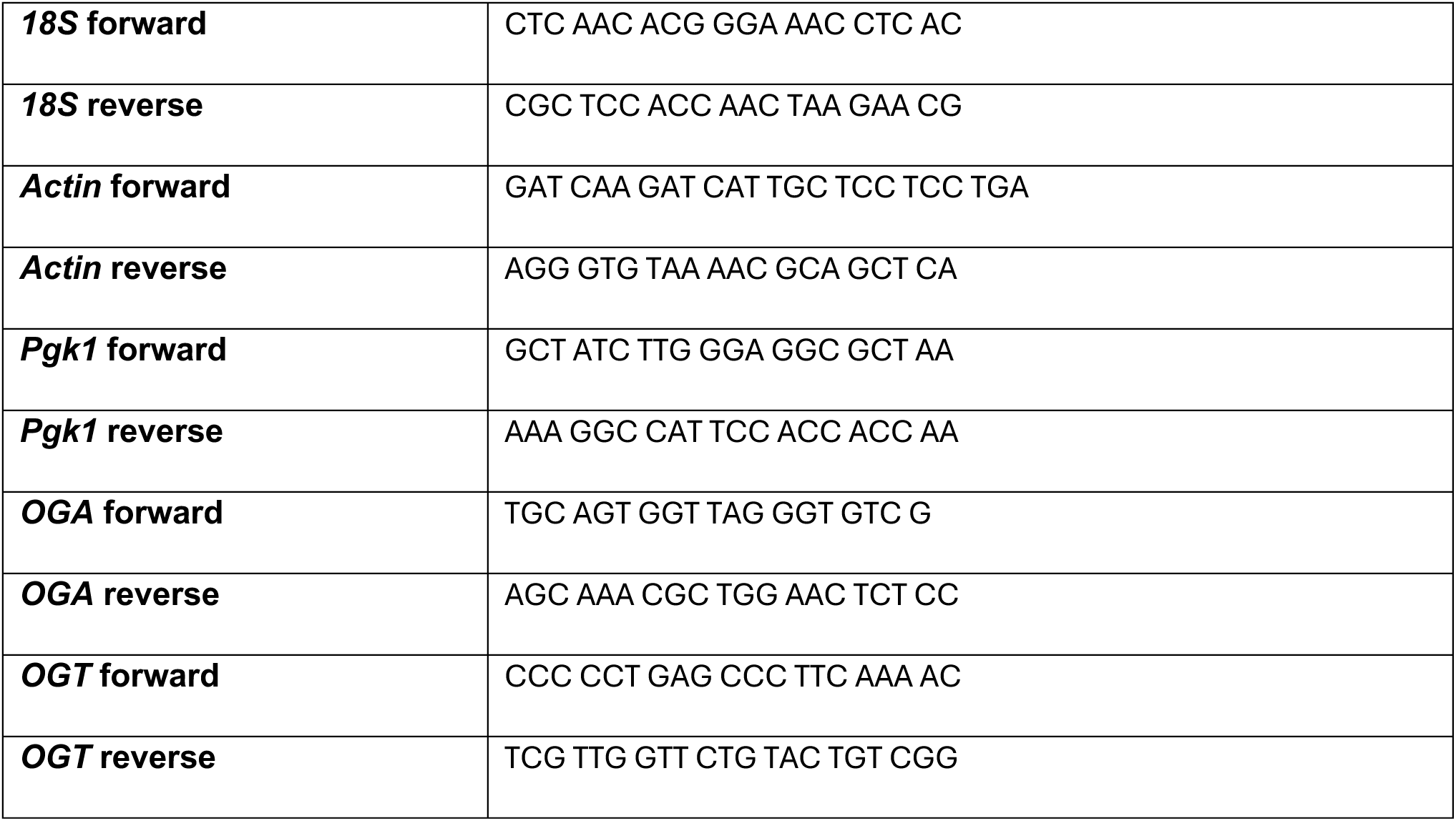
Primer sequences for qPCR.

